# An *ABCA4* loss-of-function mutation causes a canine form of Stargardt disease

**DOI:** 10.1101/329151

**Authors:** Suvi Mäkeläinen, Marta Gòdia, Minas Hellsand, Agnese Viluma, Daniela Hahn, Karim Makdoumi, Caroline J. Zeiss, Cathryn Mellersh, Sally L. Ricketts, Kristina Narfström, Finn Hallböök, Björn Ekesten, Göran Andersson, Tomas F. Bergström

**Affiliations:** Department of Animal Breeding and Genetics, Swedish University of Agricultural Sciences, Uppsala, Sweden; Department of Neuroscience, Uppsala University, Uppsala, Sweden; Department of Ophthalmology, Faculty of Medicine and Health, Örebro University, Sweden; Yale University School of Medicine, New Haven, Connecticut, United States of America; Canine Genetics Research Group, Kennel Club Genetics Centre, Animal Health Trust, Lanwades Park, Kentford, Newmarket, Suffolk, United Kingdom; Section for Comparative Ophthalmology, College of Veterinary Medicine, University of Missouri-Columbia. Missouri, United States of America; Department of Clinical Sciences, Swedish University of Agricultural Sciences, Uppsala, Sweden

## Abstract

Autosomal recessive retinal degenerative diseases cause visual impairment and blindness in humans and dogs. Currently, no standard treatment is available but pioneering gene therapy-based canine models have been instrumental for clinical trials in humans. To study a novel form of retinal degeneration in Labrador retriever dogs with clinical signs indicating cone and rod degeneration, we used whole-genome sequencing of an affected sib-pair and their unaffected parents. A frameshift insertion in the ATP binding cassette subfamily A member 4 (*ABCA4*) gene (c.4176insC), leading to a premature stop codon in exon 28 (p.F1393Lfs1395) was identified. In contrast to unaffected dogs, no full-length ABCA4 protein was detected in the retina of an affected dog. The *ABCA4* gene encodes a membrane transporter protein localized in the outer segments of rod and cone photoreceptors. In humans, the *ABCA4* gene is associated with Stargardt disease (STGD), an autosomal recessive retinal degeneration leading to central visual impairment. A hallmark of STGD is the accumulation of lipofuscin deposits in the retinal pigment epithelium. The discovery of a canine homozygous *ABCA4* loss-of-function mutation may advance the development of dog as a large animal model for human STGD.

*Author summary:* Stargardt disease (STGD) is the most common inherited retinal disease causing visual impairment and blindness in children and young adults, affecting 1 in 8-10 thousand people. For other inherited retinal diseases, the dog has become an established comparative animal model, both for identifying the underlying genetic causes and for developing new treatment methods. To date, there is no standard treatment for STGD and the mouse model is the only available animal model to study the disease. As a nocturnal animal, the morphology of the mouse eye differs from humans and therefore the mouse model is not ideal for developing methods for treatment. We have studied a novel form of retinal degeneration in Labrador retrievers showing clinical signs similar to human STGD. To investigate the genetic cause of the disease, we used whole-genome sequencing of a family quartet including two affected offspring and their unaffected parents. This led to the identification of a loss-of-function mutation in the *ABCA4* gene. The findings of this study may enable the development of a canine model for human STGD.

## Introduction

Inherited retinal dystrophies are a genetically and clinically heterogeneous group of eye diseases leading to severe visual impairment in both humans and dogs [1-7]. These diseases include various forms of retinitis pigmentosa (RP), Leber congenital amaurosis (LCA), age-related macular degeneration (AMD), cone-rod dystrophies (CRD), and Stargardt disease (STGD) and are caused by many different mutations leading to deterioration of neuroretinal and retinal pigment epithelial (RPE) function. Over 100 years ago, progressive retinal atrophy (PRA) was described as a canine equivalent of human RP [8] and is today the most common inherited retinal degenerative disease in dogs [9]. The shared phenotypic similarity of inherited retinal dystrophies in dogs and humans has made canine models attractive for gene discovery and for experimental treatments, including gene therapy, of inherited degenerative retinal disease [1, 7, 10-13]. The development of gene therapy for *RPE65*-mediated LCA is an example where a canine comparative model has been instrumental for proof-of-principle trials [10, 11, 14-16]. The identification of the p.C2Y mutation (OMIM: 610598.0001) in the *PRCD* gene is another illustrative example of the benefits of using canine genetics to find homologous candidate genes for human retinal dystrophies; the *PRCD* gene was initially mapped and identified in PRA affected dogs and subsequently in a human family with RP [17]. This mutation segregates in multiple dog breeds, including the Labrador retriever, where no other causative genetic variants for inherited retinal degenerations have been identified. In this study, a Labrador retriever sib-pair, one male and one female, negative for the p.C2Y mutation, was diagnosed with a novel form of retinal disease. To identify genetic variants associated with this novel canine retinal disease, we performed whole-genome sequencing (WGS) of the two affected individuals and their unaffected parents.

## Results and discussion

The affected dogs were visually impaired under both daylight and dim light conditions. Ophthalmoscopy revealed abnormal mottling of both central and peripheral retina, reduced reflection of light, as well as subtle retinal vascular attenuation (**Fig 1a**). A patchy outer retinal atrophy was observed with optical coherence tomography (OCT) (**Fig 1b**). In contrast, retinal layering and thickness of both outer and inner retinal layers appeared similar in an unaffected dog and a carrier (**Fig 1b**). Compared to a retina of a normal dog (**Fig 1c)**, loss of cones, less densely packed photoreceptor nuclei, increased lipofuscin accumulation in the RPE, as well as multifocal RPE hyperplasia and hypertrophy with focal atrophy of the overlying neuroretina were observed with light-microscopy in the affected male (**Fig 1d**). We used flash-electroretinography (FERG) to study the photoreceptor function in the three dogs. The inclination of the first part of the a-waves of the dark-adapted FERG in response to a bright flash was less steep and the amplitude of the a-waves were lower in both a carrier and an affected dog compared to the age-matched unaffected dog (**Fig 1e**). The a-wave of the affected dog is widened with longer implicit time. Furthermore, oscillatory potentials are less conspicuous and the first part of the b-wave is essentially lost (**Fig 1e**). Light-adapted cone transient responses (**Fig 1f**) and cone flicker (**Fig 1g**) were profoundly abnormal in the affected dog, but considerably closer to normal in the carrier. In summary, FERG demonstrated loss of cone function and abnormal rod responses, including abnormally slow dark-adaptation in the affected dog (**Fig 1e-g**). Taken together, clinical features were atypical for PRA, but showed similarities to human STGD.

**Figure 1:**
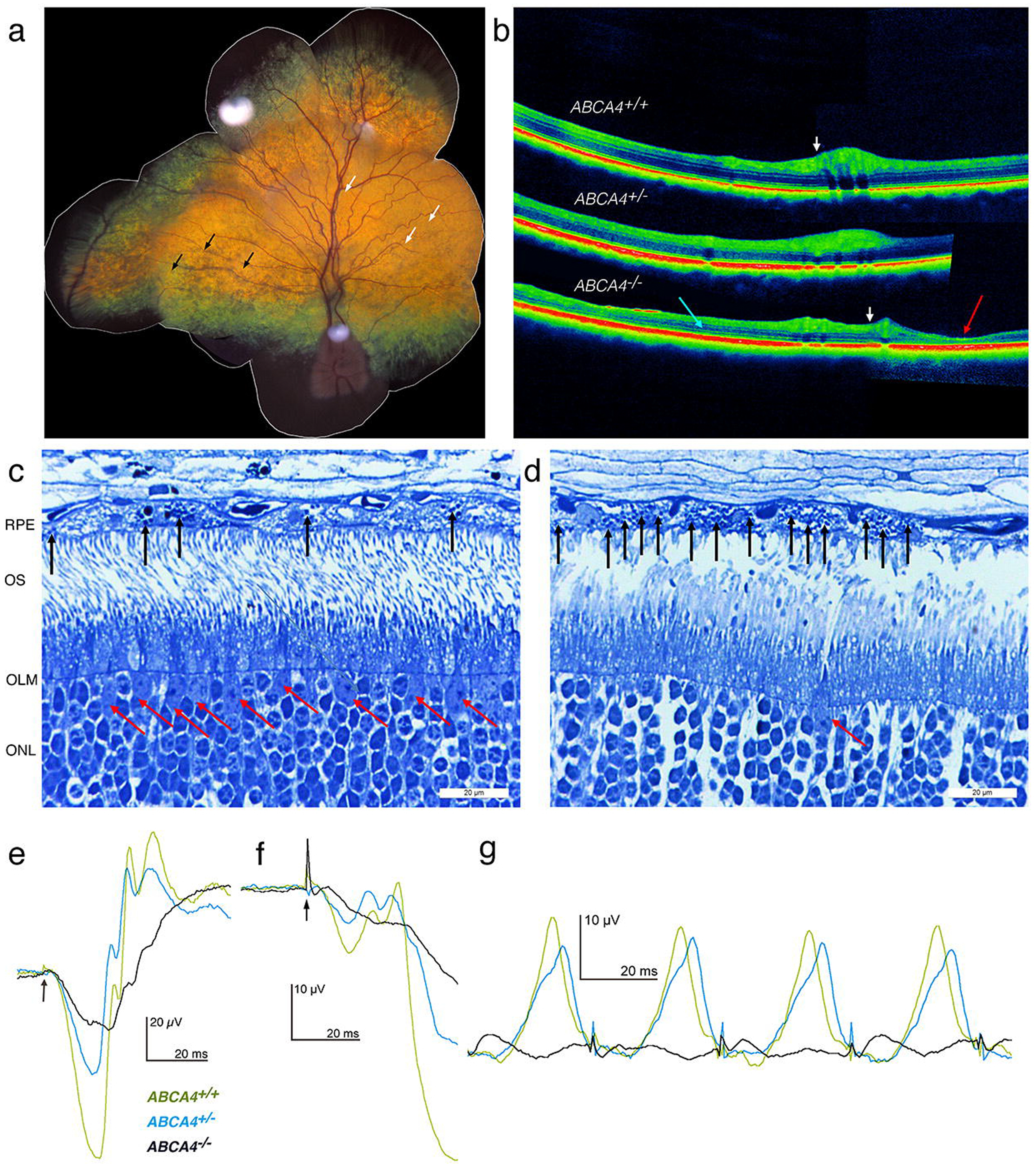
Retinal morphology and function in canine Stargardt disease. **(A)** The tapetal fundus of an affected dog. Black arrows indicate mottling (darker foci) and white arrows slight attenuation of the retinal blood vessels. **(B)** OCT images along the visual streak in age-matched unaffected dog (top), carrier (middle) and an affected dog (bottom). White arrows indicate where two images have been concatenated. A general neuroretinal thinning is visible in the affected retina (blue arrow) and includes patches of severe retinal atrophy (red arrow). **(C)** Histology of a normal canine retina and **(D)** histology of an affected retina, with red arrows indicating cone photoreceptors and black arrows indicated accumulation of lipofuscin in RPE. **(E)** Dark-adapted FERG in response to a bright flash (arrow) in an age-matched unaffected dog (green), a carrier (blue tracing) and an affected dog (black). **(F)** Light-adapted cone transient responses and **(G)** cone flicker responses with FERG. FERG = flash-electroretinography; RPE = retinal pigment epithelium; OS = outer segments; OLM = outer limiting membrane; ONL = outer nuclear layer.

The WGS of the family quartet resulted in an average coverage of 18.2x (**S1 Table**) and the identification of 6.0 × 10^6^ single nucleotide variants (SNVs) and 1.9 × 10^6^ insertions/deletions (INDELs), of which 48,299 SNVs and 5,289 INDELs were exonic. We used conditional filtering to identify 322 SNVs (of which 117 were nonsynonymous) and 21 INDELs that were consistent with an autosomal recessive pattern of inheritance (**S2 Table**). To further reduce the number of candidate variants, we compared the positions of the variants to 23 additional dog genome sequences to identify 18 nonsynonymous SNVs in 13 different genes and four INDELs in four genes that were private to the Labrador retriever family (**S2 and S3 Tables**). Fourteen of these genes were not strong candidates based on reported function and predicted effect and were not considered further. The remaining three genes, *KIAA1549*, Usherin *(USH2A),* and ATP binding cassette subfamily A member 4 (*ABCA4)* are listed in the Retinal Information Network (RetNet) database as associated with human retinal diseases and thus considered as causative candidates for canine retinal degeneration[18]. However, the variant at the *KIAA1549* gene was predicted to have a neutral effect on the protein structure (PROVEAN score −2.333, Polyphen-2 score 0.065) and was therefore discarded. The genetic variants in the *USH2A* (exon 43; c.7244C>T) and *ABCA4* (exon 28; c.4176insC) genes were validated by Sanger sequencing. Mutations in the *USH2A* gene are associated with Usher syndrome and RP, resulting in hearing loss and visual impairment [19]. The identified nonsynonymous substitution in the *USH2A* was scored as “probably damaging” using Polyphen-2 (score of 0.97) and as “deleterious” using PROVEAN (score of −4.933) (**S3 Table**). Next, we evaluated if the genetic variants of *USH2A* and *ABCA4* were concordant with the disease by genotyping eight additional clinically affected and thirteen unaffected Labradors. Out of these dogs, 16 were related to the family quartet used in the WGS (**S1 Fig**). The *USH2A* variant was discordant with the disease phenotype and was therefore excluded from further analysis (**S4 Table**). In contrast, all eight affected individuals were homozygous for the *ABCA4* insertion and the 13 unaffected individuals were either heterozygous or homozygous for the wild-type allele (**S4 Table**).

In the *ABCA4* gene, we identified a single base pair insertion of a cytosine (C) in a cytosine mononucleotide-repeat region in exon 28, where the canine reference sequence consists of seven cytosines (CanFam3.1 Chr6:55,146,549-55,146,555) (**Fig 2a**). The insertion in this region results in a non-synonymous substitution at the first codon downstream of the repeat (c.4176insC), and subsequently leads to a premature stop codon (p.F1393Lfs*1395) (**Fig 2c**). If translated, this would result in a truncation of the last 873 amino acids of the wild-type ABCA4 protein (**Fig 2b-c)**. Both the human and the dog *ABCA4* gene consists of 50 exons and encodes a ∼250 kDa ABC transporter protein (**Fig 2d**) (human and dog ABCA4 consists of 2,273 and 2,269 amino acid residues, respectively) [20, 21]. ABCA4 is localized to the disc membranes of photoreceptor outer segments and facilitates the clearance of all-*trans*-retinal from the photoreceptor discs [22-24].

**Figure 2:**
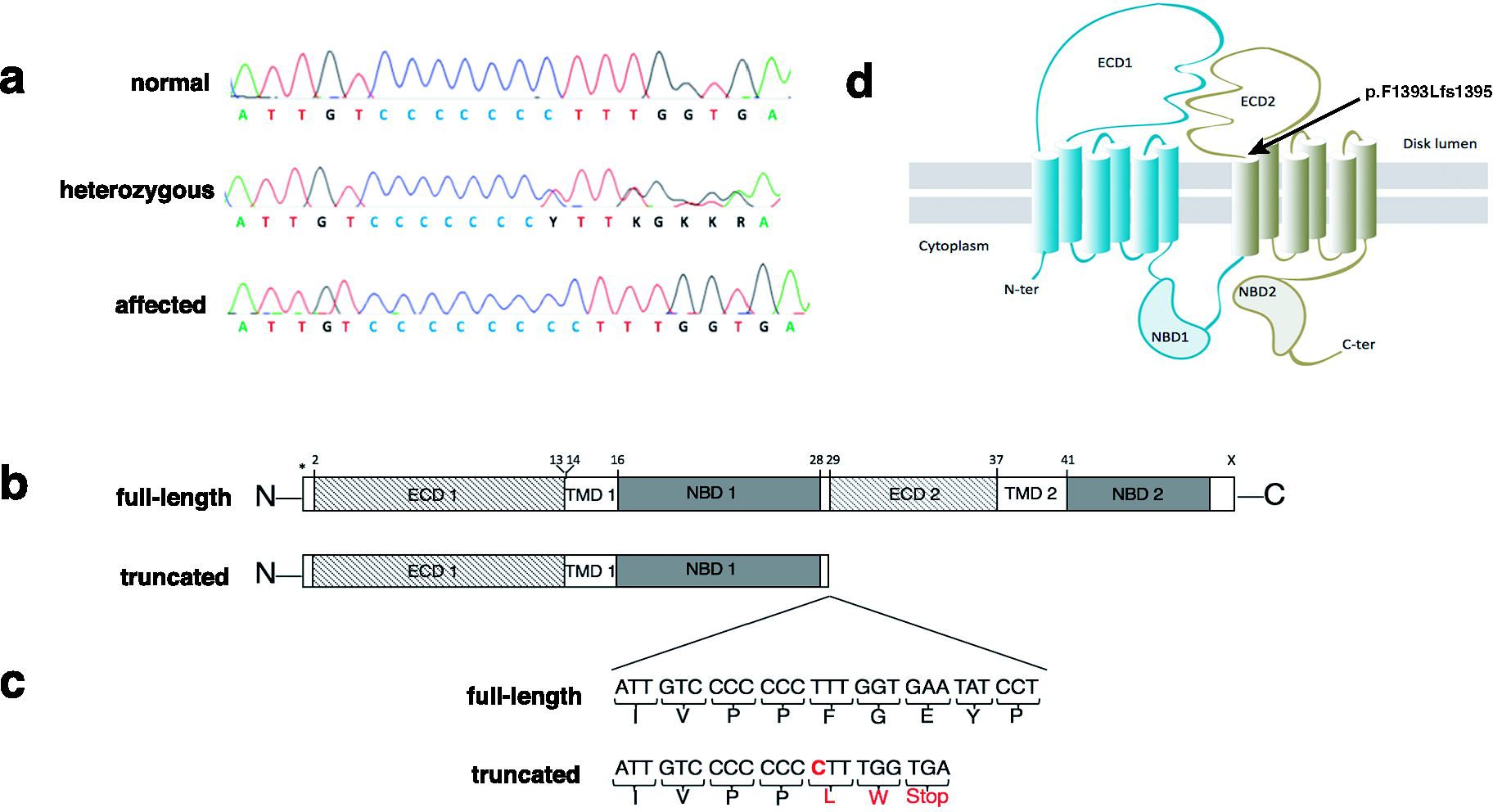
Loss-of-function mutation in the canine *ABCA4* gene. **(A)** Sanger sequencing traces spanning positions Chr6:55,146,545-55,146,564 (Canfam3.1) in exon 28 of the *ABCA4* gene of a wild-type *ABCA4*^+/+^ dog (top), a heterozygous *ABCA4*^+/-^ dog (middle), and a homozygous *ABCA4*^-/-^ dog (bottom). **(B)** Predicted structure of canine full-length ABCA4 protein, based on the proposed human structure [26], and the putative truncated product as a result of the premature stop codon at amino acid position 1,395. **(C)** Schematic representation of the region where the insertion of cytosine (C) is found showing the nucleotide and amino acid sequences of a full-length (top) and truncated (bottom) protein. **(D)** Predicted topological organization of ABCA4 and its domains with the insertion leading to a premature stop codon marked with an arrow. The topological organization is based on the proposed human topological organization [27, 28]. ECD1 = first extracellular domain; TMD1 = first membrane-spanning region; NBD1 = first nucleotide-binding domain; ECD2 = second extracellular domain; TMD2 = second membrane-spanning region; NBD2 = second nucleotide-binding domain.

To compare retinal *ABCA4* gene expression in an affected, a carrier, and a wild-type dog, we performed quantitative RT-PCR (qPCR). Primers were designed to amplify three different regions of the gene. The amplicons spanned the 5´-end (exons 2-3), the identified insertion (exons 27-28) and the 3´-end of the *ABCA4* gene (exons 47-48) (**S5 Table**). Each of the three primer pairs amplified a product of expected size in all three individuals. This suggests that despite the insertion leading to a premature stop codon in exon 28, the transcripts are correctly spliced. Relative levels of *ABCA4* mRNA were lower for the allele with the insertion in comparison to the wild-type allele (**Fig 3a**). This is consistent with nonsense-mediated decay (NMD) degrading a fraction of the transcripts with premature translation stop codon [25]. Transcripts not targeted by NMD could potentially be translated into a truncated protein of only 1,394 amino acid residues including the first extracellular domain (ECD1), the first nucleotide-binding domain (NBD1) and two membrane-spanning regions (**Fig 2b**) but lacking the second extracellular domain (ECD2) and the second nucleotide-binding domain (NBD2) [26-28] (**Fig 2b-d**). The NBDs are conserved across species and the NBD2, which is also referred to as the ATP binding cassette of the ABCA4 protein, has been shown to be particularly critical for its function as a flippase [26, 28].

**Figure 3:**
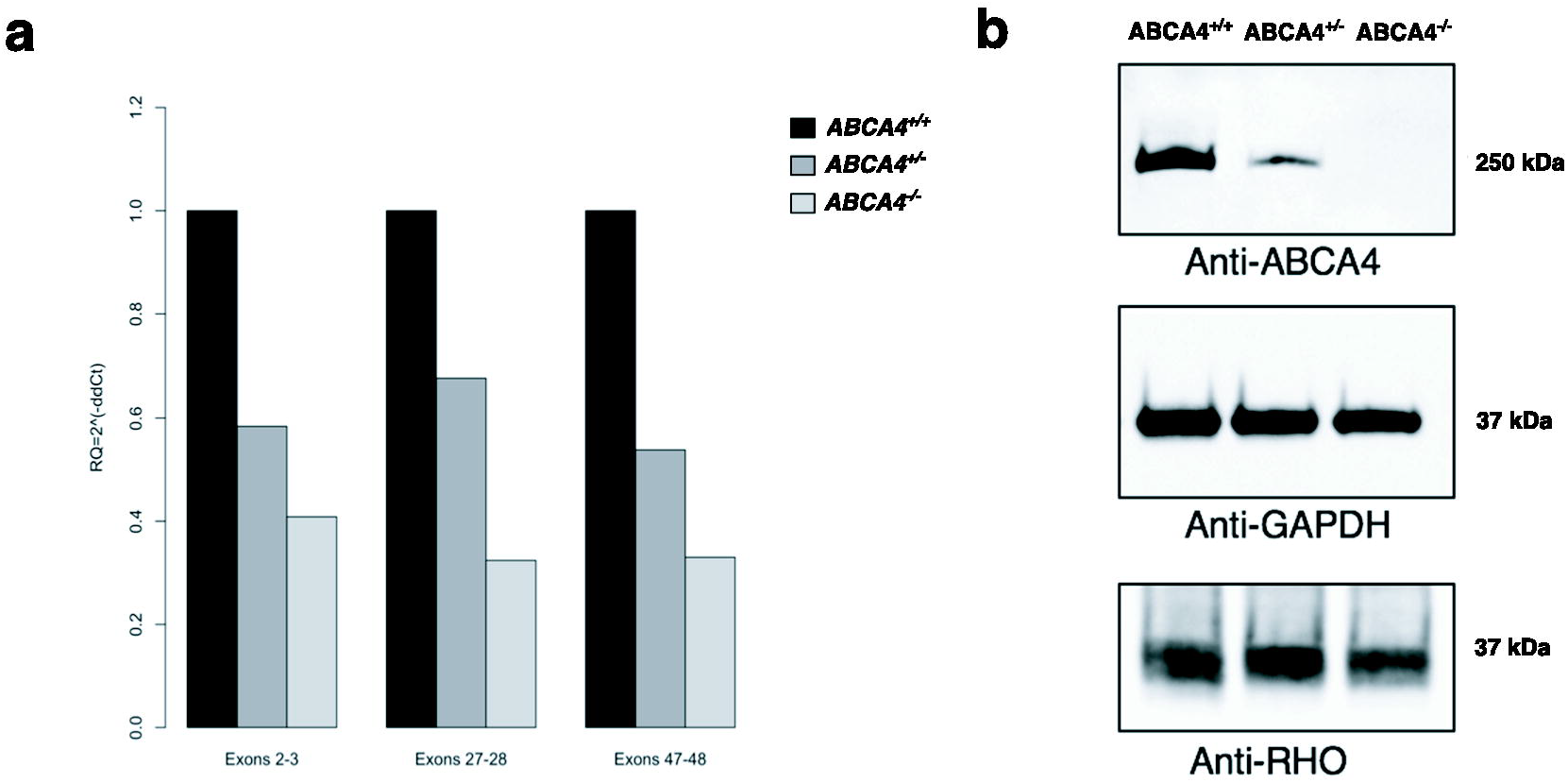
Characterization of *ABCA4* mRNA expression and western blot analyses of ABCA4 protein levels in the canine retina. **(A)** Relative *ABCA4* mRNA expression levels by quantitative RT-PCR in three different regions in three dogs with different genotypes (*ABCA4*^+/+^, *ABCA4*^+/-^ and *ABCA4*^-/-^), normalized to *GAPDH* expression. **(B)** Western blot analyses of ABCA4 (above), GAPDH (middle), and RHO (below) protein levels in retinal tissue.

To investigate the presence of full-length protein, we performed western blot analysis using an anti-ABCA4 antibody recognizing a C-terminal epitope and detecting a protein product with an approximate size of ∼250 kDa. We observed a single, correctly-sized band in samples prepared from both wild-type and heterozygous dogs. The intensity of staining in retinal protein samples from the heterozygous individual was markedly lower in comparison to the samples from the wild-type retina (**Fig 3b**). In contrast, no band was detected in the retinal sample from the affected dog. To confirm the presence of photoreceptor cells, we used an anti-RHO antibody and detected similar levels of rhodopsin in all three samples (**Fig 3b**). These results suggest that no full-length protein product is produced as a result of the insertion leading to a frameshift and a premature stop codon.

Fluorescence histochemistry was used to analyze the ABCA4 protein expression and peanut agglutinin (PNA)-binding in retinas from three dogs with different *ABCA4* genotypes. PNA binds selectively to cones in the retina [29]. ABCA4 immunoreactivity (IR) was seen in the outer part of the neural retina and the RPE. The pattern corresponded to photoreceptor outer segments and overlapped partially with the PNA label. PNA stained cone-shaped cells spanning both the inner and outer segments (**Fig 4a**). ABCA4 IR was also seen on PNA-negative outer segments, likely to be rod photoreceptors and RPE. The ABCA4 IR and PNA patterns were similar in wild-type and heterozygous retinas. In sharp contrast, no ABCA4 IR was found in the affected retina (**Fig 4a-c**). In addition, no evident PNA-staining was observed, implying loss of cones. We therefore counted photoreceptor nuclei in the three genotypes and compared the outer and inner nuclear layers. The photoreceptor nuclei are positioned in the outer nuclear layer but not in the inner nuclear layer and there were fewer nuclei in the affected outer nuclear layer in the affected retina than in the wild-type or heterozygous retina (**Fig 4d**). The corresponding reduction of nuclei was not seen in the inner nuclear layer, suggesting that photoreceptors were affected but not neurons in the inner nuclear layer. The loss of ABCA4 protein, loss of cone outer segment-PNA-staining, and the reduction of photoreceptor nuclei in the affected retina strongly imply that photoreceptors degenerate in the ABCA4^-/-^ retina.

**Figure 4:**
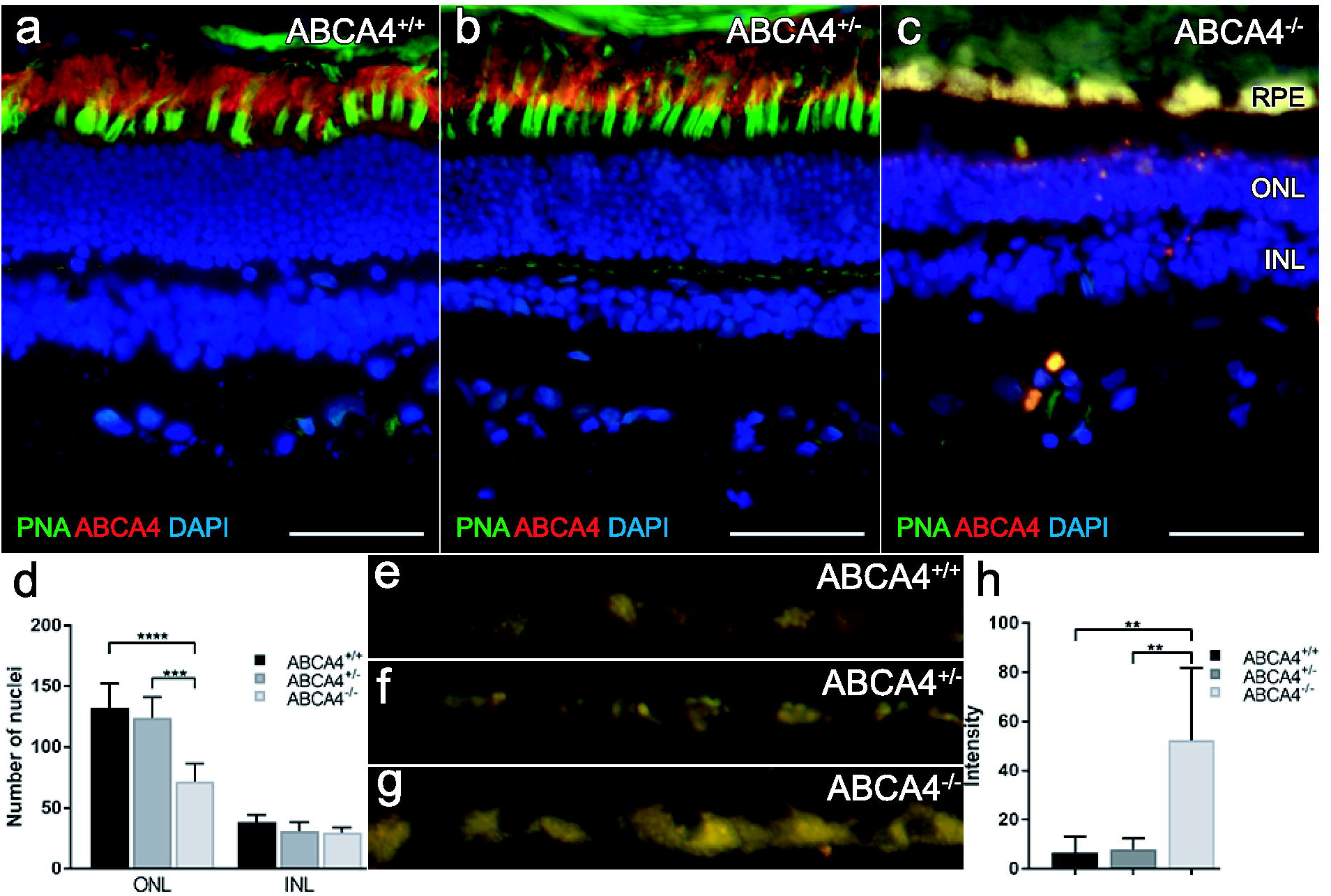
Fluorescence histochemistry of ABCA4, cone photoreceptors, and autofluorescence in the canine retina. **(A-C)** Fluorescence micrographs showing ABCA4 expression (red), FITC-conjugated peanut agglutinin (PNA, green) and DAPI nuclear staining (blue) in wild-type (ABCA4^+/+^), heterozygous (ABCA4^+/-^), and affected (ABCA4^-/-^) retinas. PNA labels cone photoreceptors. Autofluorescence, indicative of lipofuscin accumulation, was seen in the ABCA4^-/-^ RPE. **(D)** Bar graph with the average number of DAPI-stained nuclei within a given region of the ONL and the INL. **(E-G)** Fluorescence micrographs of RPE without immunohistochemistry show autofluorescence. **(H)** Bar graph with background-corrected mean autofluorescence-intensity in the RPE. Note the reduction of ABCA4-immunoreactivity, PNA binding, higher autofluorescence, and fewer nuclei in the ONL in the ABCA4^-/-^ compared to ABCA4^+/+^ or ABCA4^+/-^ retinas. All scale bars = 50 µm; RPE = retinal pigment epithelium; ONL = outer nuclear layer; INL = inner nuclear layer; Because there was only one individual per genotype, the statistics are valid for the technical replicates. ANOVA with Tukey’s post hoc test, n=6; \*\**P* < 0.01; \*\*\**P* < 0.001; \*\*\*\**P* < 0.0001; mean ± S.D.

The RPE layer of the affected retina was autofluorescent (**Fig 4c**), indicating accumulation of lipofuscin[30]. We analyzed autofluorescence in RPE from retinas of three dogs with different *ABCA4* genotypes. The autofluorescence in the affected retina was higher than in that of the retinas in the other genotypes (**Fig 4g-h)**. The higher autofluorescence indicates an increased accumulation of lipofuscin in the affected retina compared to the retinas from wild-type or heterozygous individuals.

The ABCA4 protein functions as an ATP-dependent flippase in the visual cycle, transporting *N*-retinylidene-phophatidylethanolamine (*N*-Ret-PE) from the photoreceptor disc lumen to the cytoplasmic side of the disc membrane [31, 32]. *N*-Ret-PE is a reversible adduct spontaneously formed between all-*trans*-retinal and phophatidylethanolamine (PE), and is unable to diffuse across the membrane by itself. Once transported by ABCA4, *N*-Ret-PE is dissociated and all-*trans*-retinal will re-enter to the visual cycle [33]. Defective ABCA4 leads to accumulation of *N*-Ret-PE, which together with all-*trans*-retinal, will form di-retinoid-pyridinium-phosphatidylethanolamine (A2PE) that is further hydrolyzed to phosphatidic acid (PA) and a toxic bis-retinoid, di-retinal-pyridinium-ethanolamine (A2E) [34]. This will lead to an accumulation of A2E in RPE cells when photoreceptor discs are circadially shed and phagocytosed by the RPE [35-37]. A2E is a major component of RPE lipofuscin, accounts for a substantial portion of its autofluorescence, and has a potentially toxic effect on the RPE leading to photoreceptor degeneration [35, 38-40]. In ABCA4-mediated diseases, cone photoreceptors are typically affected prior to rods [41]. The accumulation of lipofuscin and the degeneration of cones seen in this study of Labrador retrievers are also characteristic of STGD in humans [30, 42].

Mutations in the human *ABCA4* (*ABCR*) gene cause autosomal recessive STGD, autosomal recessive forms of CRD and RP [43-45]. The gene was first cloned and characterized in 1997 [20], and 873 missense and 58 loss-of-function variants have been reported in the ExAC database [46, 47], many of which are associated with visual impairment [48-50]. Currently, there is no standard treatment for STGD in humans and *Abca4*^*-/-*^ mouse is the only available animal model [51, 52]. Mice, however, lack the macula, which is primarily the area affected in STGD, and although mouse models have provided insight into genesis of the lipofuscin fluorophore A2E, *Abca4*^*-/-*^ mice do not exhibit a significant retinal degeneration phenotype [53, 54]. Unlike the mouse retina, the dog has a cone rich fovea-like area functionally similar to human fovea centralis [1, 3, 11]. The canine eye is also similar in size to the human eye, and dog has successfully been used for experimental gene therapy for retinal degenerations such as LCA, RP, rod-cone dysplasia type 1 (rcd1) [12, 16, 55]. For over a decade there has been interest in finding a canine model for *ABCA4* mediated diseases [56-58]. The loss-of-function mutation identified here can be used to develop large animal model for human STGD.

## Methods

### Animals and samples

A family quartet of Labrador retrievers (sire, dam, and two affected offspring numbered LAB1, LAB2, LAB3 and LAB4 respectively) were used in the whole-genome sequencing (WGS). In addition, 16 related individuals (LAB5 to LAB20, see **S1 Figure**) as well as five unrelated Labradors (LAB 21 to LAB25) were used to validate the WGS findings. Whole blood samples from these dogs were collected in EDTA tubes and genomic DNA was extracted using 1 ml blood on a QIAsymphony SP instrument and the QIAsymphony DSP DNA Kit (Qiagen). We obtained eyes from the affected male (LAB4) and his unaffected sibling (LAB6) at the age of 12, as well as from one unrelated, unaffected 11-year-old female Labrador retriever (LAB24) and one 10-year-old male German spaniel (GS) after their euthanization (with sodium pentobarbithal (Pentobarbithal 100 mg/ml, Apoteket Produktion & Laboratorier AB) for unrelated reasons. All samples were obtained with informed dog owner consent. Ethical approval was granted by the regional animal ethics committee (Uppsala djursförsöksetiska nämd; Dnr C12/15).

### Ophthalmic exam and optical coherence tomography (OCT)

Ophthalmic examination included reflex testing, testing of vision with falling cotton balls under dim and daylight conditions, indirect ophthalmoscopy and slit-lamp biomicroscopy. The affected male (LAB4), his unaffected sibling (LAB6) and an unaffected, age-matched, female Labrador (LAB22) were examined with spectral-domain OCT (Topcon 3D OCT-2000, Topcon Corp.). The examination was done after pupillary dilation, but without sedation, using repeated horizontal single line scans (6 mm, 1024 A-scans) (Topcon 3D OCT-2000, Topcon Corp.) along the visual streak area.

### Flash-electroretinography (FERG)

We recorded FERG bilaterally from the three dogs examined with OCT under general anaesthesia. Sedation with intramuscular acepromazine 0.03 mg/kg (Plegicil vet., Pharmaxim Sweden AB) was followed by induction with propofol 10 mg/kg IV (Propovet, Orion Pharma Animal Health AB). After intubation, inhalation anaesthesia was maintained with isoflurane (Isoflo vet., Orion Pharma Animal Health AB). Corneal electrodes (ERG-JET, Cephalon A/S) were used with isotonic eye drops (Comfort Shield, i.com medical GmbH) as coupling agent. Gold plated, cutaneous electrodes served as ground and reference electrodes (Grass, Natus Neurology Inc.) at the vertex and approximately 3 cm caudal to the lateral canthi, respectively. Light stimulation, calibration of lights and processing of signals were performed as described by Karlstam et al., 2011[59]. We used a slightly modified ECVO protocol [60], where the process of dark-adaptation was monitored for 1 hour before a dark-adapted response intensity series was performed.

### Histopathology

Sectioned eyes from the affected male (LAB4) and the unaffected male GS were immersed in Davidson’s Solution. The eyes were dehydrated in ethanol, paraffin embedded, cut into 4 µm thick sections and stained with haematoxylin and eosin (H&E).

### Whole-genome sequencing

Genomic DNA from four Labrador retriever dogs (LAB1, LAB2, LAB3 and LAB4) was fragmented using the Covaris M220 instrument (Covaris Inc.), according to the manufacturer’s instructions. To obtain sufficient sequence depth, we constructed two biological replicates of libraries with insert sizes of 350 bp and 550 bp following TruSeq DNA PCR-Free Library Prep protocol. The libraries were multiplexed and sequenced on a NextSeq500 instrument (Illumina) for 100 × 2 and 150 × 2 cycles using the High Output Kit and High Output Kit v2, respectively. The raw base calls were de-multiplexed and converted to fastq files using bcl2fastq v.2.15.0 (Illumina). The two sequencing runs from each individual were merged, trimmed for adapters and low-quality bases using Trimmomatic v.0.32[61], and aligned to the canine reference genome CanFam3.1 using Burrows-Wheeler Aligner (BWA) v.0.7.8[62]. Aligned reads were sorted and indexed using Samtools v.1.3[63] and duplicates were marked using Picard v.2.0.1. The BAM files were realigned and recalibrated with GATK v.3.7[64]. Multi-sample variant calling was done following GATK Best Practices [65] using publicly available genetic variation Ensembl Variation Release 88 in dogs (*Canis lupus familiaris*). We filtered the variants found by GATK using the default values defining two groups of analyses: trio 1 and 2, both consisting of the same sire and dam, and one of their affected offspring. Variants annotated in the exonic region with ANNOVAR v.2017.07.16 [66], presenting an autosomal recessive inheritance pattern and shared between the two trios were selected for further evaluation. To predict the effects of amino acid changes on protein function, we evaluated SNVs using PolyPhen-2 v2.2.2r398 [67] and PROVEAN v.1.1.3 [68] and nonframeshift INDELS using PROVEAN v.1.1.3. Frameshift INDELs were manually inspected using The Integrative Genomics Viewer (IGV) [69, 70]. The sequence data were submitted to the European Nucleotide Archive with the accession number PRJEB26319.

### Validation of the variants

To validate the WGS results, we designed primers amplifying the variants c.7244C>T in *USH2A* gene and c.4176insC in *ABCA4* gene with Primer3 [71, 72] (**S5 Table**) and sequenced the family quartet using Applied Biosystems 3500 Series Genetic Analyzer (Applied Biosystems, Thermo Fisher Scientific). To test if the variants were concordant with the disease, 21 additional ophthalmologically evaluated Labrador retrievers were genotyped by Sanger sequencing. Eight of these dogs were clinically affected and thirteen showed no signs of retinal degeneration by the age of seven years.

### Quantitative RT-PCR (qPCR)

Neuroretinal samples were collected from the affected dog (LAB4), the heterozygous sibling (LAB6), and the unaffected female (LAB24). The samples were immediately preserved in RNAlater (SigmaAldrich), homogenized with Precellys homogenizer (Bertin Instruments) and total RNA was extracted with RNAeasy mini kit (Qiagen) according to the manufacturer’s instructions. RNA integrity and quality were inspected with Agilent 6000 RNA Nano kit with the Agilent 2100 Bioanalyzer system (Agilent Technologies). cDNA was synthesized using RT^2^ First Strand kit (Qiagen) with random hexamers provided in the kit. cDNA concentration was inspected with Qubit ssDNA Assay kit (Life Technologies). RT^2^ qPCR Primer Assay (Qiagen) was used to amplify the reference gene *GAPDH*. To amplify the target gene *ABCA4*, we designed custom primers with Primer3 [71, 72] targeting three different regions spanning exons 2 to 3, 27 to 28, and 47 to 48 (**S5 Table**). We amplified the cDNA fragments encoding regions of interest using RT^2^ SYBR Green ROX qPCR Mastermix (Qiagen) with StepOnePlus Real-Time PCR system (Applied Biosystems, Thermo Fisher Scientific), according to the manufacturer’s instructions. Target gene expression was normalized to expression of *GAPDH*, and shown relative to a control *ABCA4*^+/+^ sample (ΔΔC_T_ method). The results were confirmed in two independent experiments.

### SDS-Gel Electrophoresis and Western Blotting

We extracted protein from the neuroretinal samples of the individuals used in qPCR (see above) by homogenization in Pierce RIPA lysis buffer (Thermo Scientific) supplemented with phosphatase inhibitor cocktail (Sigma, P8340) using the Precellys homogenizer (Bertin Instruments). Protein concentration was determined using the Pierce BSA Protein Assay kit (Thermo Fischer Scientific). 50 μg of protein samples were resolved by SDS-PAGE, transferred to nitrocellulose membrane, and immunoblotted with the following primary antibodies: ABCA4 (Novus Biologicals, NBP1-30032, 1:1000), GAPDH (Thermo Scientific, MA5-15738, 1:1000), Rhodopsin (Novus Biologicals, NBP2-25160SS, 1:5000), followed by Anti-Mouse IgG horseradish peroxidase-conjugated secondary antibody (R&D Systems, HAF007, 1:5000). Binding was detected using the Clarity western ECL substrate (Bio-Rad).

### Fluorescence histochemistry

Tapetal fundus from the affected male (LAB4), his heterozygous sibling (LAB6), and the unaffected GS were fixed in 4% PFA in 1x PBS on ice for 15 minutes, washed in 1x PBS for 10 minutes on ice, and cryoprotected in 30% sucrose overnight at 4°C. The central part of the fundus was embedded in Neg-50™ frozen section medium (Thermo Scientific), and 10 µm sections were collected on Superfrost Plus slides (J1800AMNZ, Menzel-Gläser). The sections were re-hydrated in 1x PBS for 10 minutes, incubated in blocking solution (1% donkey serum, 0.02% thimerosal, and 0.1% Triton X-100 in 1x PBS) for 30 minutes at room temperature, and incubated in primary antibody ABCA4 (1:500, NBP1-30032, Novus Biologicals) and FITC-conjugated lectin PNA (1:400, L21409, Molecular Probes) solution at 4°C overnight. Following overnight incubation, the slides were washed 3 × 5 minutes in 1x PBS and incubated in Alexa 568 secondary antibody (1:2000, A10037, Invitrogen) solution for at least 2 hours at room temperature and washed 3 × 5 minutes in 1x PBS. The slides were mounted using ProLong^®^ Gold Antifade Mountant with DAPI (P36931, Molecular Probes). Fluorescence images were captured using a Zeiss Axioplan 2 microscope equipped with an AxioCam HRc camera.

### Counting nuclei

Ten micrometer retinal sections were mounted as described under Fluorescence histochemistry, and the number of nuclei within a region with a width of 67 µm that was perpendicular to and covered both the outer and inner nuclear layers were counted. Nuclei in the outer and inner nuclear layers were counted separately. We analysed six images from each of the three animals (LAB4, LAB6, and GS). Bar graphs were generated and statistical analysis of the technical replicates (one-way ANOVA with Tukey’s post hoc multiple comparison analysis) was performed in GraphPad Prism 7.

### Autofluorescence

Retinal sections were washed, incubated in blocking solution, and mounted as described under Fluorescence histochemistry. The exposure times for the excitation at 488 nm and 568 nm were fixed for all images taken (150 ms and 80 ms, respectively). Outlines of the retinal pigment epithelium (RPE), as well as adjacent background regions, were drawn using the polygon selection tool in ImageJ (v1.51, NIH), and the area and mean fluorescence intensity were measured. The mean intensity of the autofluorescence in the RPE was calculated by subtracting the background intensity from the adjacent regions. We analysed six images from each of the three individuals used in the fluorescence histochemistry. Bar graph generation and statistical analysis were performed as described under Counting nuclei.

## Acknowledgements

The authors would like to thank the veterinary clinicians Berit Wallin-Håkansson, Ida Möller and Stuart Ellis for diagnosing and collecting samples, Anna Svensson and Kiran Kumar Jagarlamudi for technical assistance with western blotting, Mihaela Martis at the Swedish Bioinformatics Infrastructure Sweden at SciLifeLab for bioinformatics advice and Kerstin Lindblad-Toh for valuable comments of earlier versions of the manuscript. We would also like to acknowledge the support of the dedicated dog owners who allowed their dogs to take part in this study. The authors would like to acknowledge the support of the National Genomics Infrastructure (NGI) / Uppsala Genome Center and UPPMAX for providing computational infrastructure.

## Author contributions

G.A. and T.B. conceived, designed and directed the study; B.E. and K.N. collected field material and diagnosed the subjects; B.E. and K.M. performed OCT, C.J.Z. performed histological analysis; C.M. and S.R. provided samples and genotypes; M.G., S.M. and D.H. performed sequencing; S.M. and M.G. performed sequence analyses and bioinformatics with support from A.V.; S.M. performed qPCR and western blotting; M.H. and F.H. performed and analyzed fluorescence histochemistry experiments; S.M., T.B. and G.A. were responsible for preparing of the manuscript with particular contributions from M.G., M.H., F.H., B.E. and C.M; all authors read and approved the final manuscript.

## Supporting information

**S1 Fig. Pedigree of the Labrador retriever dogs used in the study.** Filled symbols indicate affected individuals, half-filled symbols represent obligate or genotyped carriers of the *ABCA4* insertion. Individuals LAB1 to LAB4 were used in the WGS analysis. Numbered individuals were genotyped for the insertion in the *ABCA4* gene (c.4176insC) and for the non-synonymous substitution in the *USH2A* gene (c.7244C>T). In addition, five unrelated, unaffected dogs (not shown in the figure) were genotyped and found to be either wild-type or heterozygous for the variants in the *ABCA4* and the *USH2A* gene (LAB21 to LAB25).Crosses intersecting the dashed lines indicate the number of generations between the individuals.

**S1 Table. Summary of the whole-genome sequencing runs 1 and 2.**

**S2 Table. Exonic variants identified in WGS.** Number of exonic variants following autosomal recessive inheritance pattern (AR) in Trio1 and Trio2, each consisting of the parents and one of the two offspring. The total number of exonic variants in the family quartet including all inheritance patterns and the number of AR variants shared between the two trios. The “unique” column represents the number of AR variants, which were shared between the two trios and not found to be homozygous in 23 additional investigated canine genome sequences.

**S3 Table. List of candidate variants from WGS.** Coding sequence variants identified as private for the Labrador retriever family and the predicted effect of the variants based on Polyphen-2 and PROVEAN scores.

**S4 Table. Validation of variants c.4176insC in ABCA4 gene and c.C7244T in USH2A gene by Sanger sequencing.**

**S5 Table. Canine primer sequences used in the analysis.**

